# General DNA methylation patterns and environmentally-induced differential methylation in the eastern oyster (*Crassostrea virginica*)

**DOI:** 10.1101/2020.01.07.897934

**Authors:** Yaamini R. Venkataraman, Alan M. Downey-Wall, Justin Ries, Isaac Westfield, Samuel J. White, Steven B. Roberts, Kathleen E. Lotterhos

## Abstract

Epigenetic modification, specifically DNA methylation, is one possible mechanism for intergenerational plasticity. Before inheritance of methylation patterns can be characterized, we need a better understanding of how environmental change modifies the parental epigenome. To examine the influence of experimental ocean acidification on eastern oyster (*Crassostrea virginica*) gonad tissue, oysters were cultured in the laboratory under control (491 ± 49 μatm) or high (2550 ± 211 μatm) *p*CO_2_ conditions for four weeks. DNA from reproductive tissue was isolated from five oysters per treatment, then subjected to bisulfite treatment and DNA sequencing. Irrespective of treatment, DNA methylation was primarily found in gene bodies with approximately 22% of CpGs (2.7% of total cytosines) in the *C. virginica* genome predicted to be methylated. In response to elevated *p*CO_2_, we found 598 differentially methylated loci primarily overlapping with gene bodies. A majority of differentially methylated loci were in exons (61.5%) with less intron overlap (31.9%). While there was no evidence of a significant tendency for the genes with differentially methylated loci to be associated with distinct biological processes, the concentration of these loci in gene bodies, including genes involved in protein ubiquitination and biomineralization suggests DNA methylation may be important for transcriptional control in response to ocean acidification. Changes in gonad methylation also indicate potential for these methylation patterns to be inherited by offspring. Understanding how experimental ocean acidification conditions modify the oyster epigenome, and if these modifications are inherited, allows for a better understanding of how ecosystems will respond to environmental change.

## Introduction

As increased anthropogenic carbon dioxide is expected to create adverse conditions for calcifying organisms (IPCC 2019), efforts have been made to understand how ocean acidification impacts ecologically and economically important organisms like bivalves (Parker et al., 2013; Ekstrom et al., 2015). Bivalve species are sensitive to reduced aragonite saturation associated with ocean acidification, with larvae being particularly vulnerable (Barton et al., 2012; Waldbusser et al., 2014). Shell structure may be compromised in larvae, juveniles, and adults (Gazeau et al., 2007; Kurihara et al., 2007; Beniash et al., 2010; Ries, 2011). Aside from affecting calcification and shell growth, ocean acidification can impact protein synthesis, energy production, metabolism, antioxidant responses, and reproduction (Tomanek et al., 2011; Timmins-Schiffman et al., 2014; Dineshram et al., 2016; Boulais et al., 2017; Omoregie et al., 2019).

Additionally, adult exposure to ocean acidification may impact their larvae (reviewed in (Ross et al., 2016; Byrne et al., 2019). For example, adult Manila clams (*Ruditapes philippinarum*) and mussels (*Musculista senhousia*) reproductively conditioned in high *p*CO_2_ waters yield offspring that exhibit significantly faster development or lower oxidative stress protein activity in those same conditions (Zhao et al., 2018, 2019). In contrast, northern quahog (hard clam; *Mercenaria mercenaria*) and bay scallop (*Argopecten irradians*) larvae may be more vulnerable to ocean acidification and additional stressors when parents are reproductively conditioned in high *p*CO_2_ waters (Griffith and Gobler, 2017). Some species exhibit both positive and negative carryover effects (ex. *Saccostrea glomerata*; (Parker et al., 2012, 2017). Intergenerational effects have also been documented when adult exposure to ocean acidification does not coincide with reproductive maturity (*e.g*. *Crassostrea gigas*; (Venkataraman et al., 2019)). Although intergenerational carryover effects are now at the forefront of ocean acidification research in bivalve species, the mechanisms responsible for these effects are still unclear.

Epigenetics is the next frontier for understanding how environmental memory may modulate phenotypic plasticity across generations (Eirin-Lopez and Putnam, 2018). Epigenetics refers to changes in gene expression that do not arise from changes in the DNA sequence, with methylation of cytosine bases being the most studied mechanism (Bird, 2002; Deans and Maggert, 2015). Unlike highly methylated vertebrate genomes, marine invertebrate taxa have sparse methylation throughout their genomes, similar to a mosaic pattern (Suzuki and Bird, 2008). Genes that benefit from stable transcription, such as housekeeping genes, tend to be more methylated, while environmental response genes that are less methylated are prone to more spurious transcription and alternative splicing patterns, thereby possibly increasing phenotypic plasticity (Roberts and Gavery, 2012; Dimond and Roberts, 2016; Gatzmann et al., 2018). Increased levels of DNA methylation can also correlate with increased transcription. Several base pair resolution studies in *C. gigas* demonstrate a positive association between DNA methylation and gene expression that is consistent across cell types (Roberts and Gavery, 2012; Gavery and Roberts, 2013; Olson and Roberts, 2014). Since DNA methylation could provide a direct link between environmental conditions and phenotypic plasticity via influencing gene activity, elucidating how invertebrate methylomes respond to abiotic factors is crucial for understanding potential acclimatization mechanisms (Bossdorf et al., 2008; Hofmann, 2017).

While bivalve species have been used as model organisms to characterize marine invertebrate methylomes, how ocean acidification affects bivalve DNA methylation is poorly understood. Methylation responses to ocean acidification have been studied in multiple coral species. When placed in low pH conditions (7.6-7.35), *Montipora capitata* did not demonstrate any differences in calcification, metabolic profiles, or DNA methylation in comparison to clonal fragments in ambient pH (7.9-7.65) (Putnam et al., 2016). DNA methylation increased in another coral species, *Pocillopora damicornis*, in addition to reduced in calcification and more differences in metabolic profiles (Putnam et al., 2016). The coral *Stylophora pistillata* also demonstrates increased global methylation as pH decreases (pH_treatment_ = 7.2, 7.4, 7.8; pH_control_ = 8.0), with methylation reducing spurious transcription (Liew et al., 2018b). Combined whole genome bisulfite sequencing and RNA sequencing revealed differential methylation and expression of growth and stress response pathways controlled differences in cell and polyp size between treatments (Liew et al., 2018b). The association between DNA methylation and phenotypic differences in these corals demonstrates that epigenetic regulation of genes is potentially important for acclimatization and adaptation to environmental perturbation. Recent examination of *C. virginica* methylation patterns in response to a natural salinity gradient suggests that differential methylation may modulate environmental response in this species (Johnson and Kelly, 2019).

There is evidence that suggests that methylation patterns can be inherited in marine invertebrates. For example, purple sea urchin (*Strongylocentrotus purpuratus*) offspring have methylomes that reflect maternal rearing conditions (Strader et al., 2019). Different parental temperature and salinity regimes influence larval methylomes in *Platygyra daedalea* (Liew et al., 2018a). In the Pacific oyster (*C. gigas*), parental exposure to pesticides influence DNA methylation in spat, even though the spat were not exposed to these conditions (Rondon et al., 2017). Methylation changes in gametes are likely the ones that could be inherited, and may play a role in carryover effects. Before determining if DNA methylation is a viable mechanism for altering the phenotypes of offspring or subsequent generations, the epigenome of bivalve reproductive tissue in response to ocean acidification must be characterized.

The present study is the first to determine if ocean acidification induces differential methylation in reproductive tissue in the eastern oyster (*Crassostrea virginica*). Adult *C. virginica* were exposed to control or elevated *p*CO_2_ conditions. We hypothesize that ocean acidification will induce differential methylation in *C. virginica* gonad tissue, and that genes with differentially methylated loci will have biological functions that could allow for acclimatization to environmental perturbation. Understanding how experimental ocean acidification conditions modify the oyster epigenome, and if these modifications are inherited, allows for a better understanding of how ecosystems will respond to environmental change.

## Methods

### Experimental Design

Adult *C. virginica* (9.55 cm ± 0.45) were collected from an intertidal oyster reef in Plum Island Sound, MA (42.681764, −70.813498) in mid-July 2016. The oysters were transported to the Marine Science Center at Northeastern University (Nahant, MA), where they were cleaned and randomly assigned to one of six flow-through tanks (50L) maintained at ambient seawater conditions. Oysters were acclimated for 14 days under control conditions (500 μatm; 14-15°C) before initiating a 28-day experimental exposure. Half of the tanks remained at control *p*CO_2_ conditions (500 μatm, Ω_calcite_ > 1), while the other half were ramped up to elevated *p*CO_2_ conditions (2500 μatm, Ω_calcite_ < 1) over 24 hours. This elevated treatment is consistent with observations in other estuarine ecosystems that oysters inhabit (Feely et al., 2010), although pH in nature only stays as extreme for short periods of time (e.g. hours). Moreover, the extreme treatment was also chosen to increase precision and therefore power to detect a response (Whitlock and Schluter, 2014).

Treatment conditions were replicated across three tanks, with oysters distributed evenly among tanks (1-2 oysters per tank). Each tank had an independent flow-regulator that delivered fresh, natural seawater at approximately 150 ml min^-1^. Carbonate chemistry was maintained independently for each tank by mixtures of compressed CO_2_ and compressed air at flow rates proportional to the target *p*CO_2_ conditions. Gas flow rates were maintained with *Aalborg* digital solenoid-valve-controlled mass flow controllers (Model GFC17, precision = 0.1mL/min). Within a treatment, tanks were replenished with fresh seawater and each tank was independently bubbled with its own mixed gas stream, with partial recirculation and filtration with other tanks in the treatment. As a result, the carbonate chemistry (i.e., the independent variable by which the treatments were differentiated) of the replicate tanks were slightly different from each other, which is evidence of their technical independence. Temperature was maintained at 15°C using Aqua Euro USA model MC-1/4HP chillers coupled with 50-watt electric heaters. Average salinity was determined by the incoming natural seawater and reflected ambient ocean salinity of Massachusetts Bay near the Marine Science Center (Latitude = 42.416100, Longitude = − 70.907737).

Oysters were fed 2.81 mL/day of a 10% Shellfish Diet 1800 twice daily following Food and Agriculture Organization’s best practices for oysters (Helm and Bourne, 2004). Five oysters were collected from each treatment at the end of the 28 day exposure. They were immediately dissected with gonadal tissue harvested and immediately flash frozen. Partial gamete maturation was evident upon visual inspection.

### Measurement and control of seawater carbonate chemistry

The carbonate chemistry of tanks was controlled by bubbling mixtures of compressed CO_2_ and compressed air at flow rates proportional to the target *p*CO_2_ conditions. The control *p*CO_2_ treatments were maintained by bubbling compressed ambient air only.

Temperature, pH, and salinity of all replicate tanks was measured three times per week for the duration of the experiment. Temperature was measured using a glass thermometer to 0.1°C accuracy, pH was measured using an *Accumet* solid state pH electrode (precision = 1mV), salinity was measured using a *YSI 3200* conductivity probe (precision = 0.1 ppt). Every two weeks, seawater samples were collected from each replicate tank for analysis of dissolved inorganic carbon (DIC) and total alkalinity (A_T_). Samples were collected in 250 mL borosilicate glass bottles sealed with a greased stopper, immediately poisoned with 100 μL saturated HgCl_2_ solution, and then refrigerated. Samples were analyzed for DIC via coulometry and Alk_T_ via closed-cell potentiometric Gran Titration with a VINDTA 3C (Marianda Corporation). Other carbonate system parameters, including Ω_calcite_, pH, and *p*CO_2_, were calculated from DIC, A_T_, salinity, and temperature using CO2SYS software version 2.1 (Lewis and Wallace, 1998; Van Heuven et al., 2011), using the seawater pH scale (mol/kg-SW) with K1 and K2 values from (Roy et al., 1993), a KHSO_4_ value from (Dickson, 1990), and a [B]_T_ value from (Lee et al., 2010).

### MBD-BS Library Preparation

DNA was isolated from five gonad tissue samples per treatment using the E.Z.N.A. Mollusc Kit (Omega) according to the manufacturer’s instructions. Isolated DNA was quantified using a Qubit dsDNA BR Kit (Invitrogen). DNA samples, ranging from 12.8 ng/μL to 157 ng/μL, were placed in 1.5 mL centrifuge tubes and sonicated using a QSONICA CD0004054245 (Newtown, CT) in 30 second interval periods over ten minutes at 4 °C and 25% intensity. Shearing size (350bp) was verified using a 2200 TapeStation System (Agilent Technologies). Samples were enriched for methylated DNA with the MethylMiner kit (Invitrogen). A single-fraction elution using 400 μL of high salt buffer was used to obtain captured DNA. After ethanol precipitation, 25 μL of buffer was used for the final elution. Library preparation and sequencing was performed by ZymoResearch using Pico Methyl-Seq Library Prep Kit (Cat. #D5455). Libraries were then barcoded and pooled into two lanes (eight samples in one and two in another) to generate 100bp paired-end reads on the HiSeq1500 sequencer (Illumina, Inc.).

### Global Methylation Characterization

Sequences were trimmed with 10 bp removed from both the 5’ and 3’ ends using TrimGalore! v.0.4.5 (Martin, 2011). Quality of sequences was assessed with FastQC v.0.11.7 (Andrews, 2010). The *C. virginica* genome (NCBI Accession GCA_002022765.4) was prepared using Bowtie 2-2.3.4 (Linux x84_64 version; (Langmead and Salzberg, 2012)) within the `bismark_genome_preparation` function in Bismark v.0.19.0 (Krueger and Andrews, 2011). Trimmed sample sequences were then aligned to the genome using Bismark v.0.19.0 (Krueger and Andrews, 2011) with non-directionality specified alignment score set using `-score_min L,0,-1.2`. Alignment files (ie. bam) were deduplicated (`deduplicate_bismark`), sorted and indexed using SAMtools v.1.9 (Li et al., 2009). Methylation calls were extracted from deduplicated files using `bismark_methylation_extractor`.

Various *C. virginica* genome feature tracks were created for downstream analyses using BEDtools v2.26.0 (Quinlan and Hall, 2010). Genes, mRNA, coding sequences, and exons were derived directly from the *C. virginica* genome on NCBI (Gómez-Chiarri et al., 2015). The complement of the exon track was used to identify introns, and coding sequences were subtracted from exons to identify untranslated regions of exons (UTR). Exon locations were removed from the complement of the gene track to define intergenic regions. Putative promoter regions were defined as those 1kb upstream of transcription start sites. Putative transposable elements were identified using RepeatMasker (v4.07) with RepBase-20170127 and RMBlast 2.6.0 (Smit et al., 2013; Bao et al., 2015). All species available in RepBase-20170127 were used to identify transposable elements.

Overall *C. virginica* gonad methylation patterns were characterized using information from all samples. Individual CpG dinucleotides with at least 5x coverage in each sample were classified as methylated (≥ 50% methylation), sparsely methylated (10-50% methylation), or unmethylated (< 10% methylation). The locations of all methylated CpGs were characterized in relation to putative promoter regions, UTR, exons, introns, transposable elements, and intergenic regions. We tested the null hypothesis that there was no association between the genomic location of CpG loci and methylation status (all CpGs versus methylated CpGs) with a chi-squared contingency test (`chisq.test` in R Version 3.5.0).

Methylation islands were determined to characterize overall methylation in the *C. virginica* genome using a sliding window analysis based on (Jeong et al., 2018). Islands were defined as areas of the genome with enriched levels of methylated CpGs (>50% methylation). To define methylation islands, each chromosome was examined using an initial 500 bp window starting at the first methylated CpG. If the proportion of methylated CpGs in the window was greater than 0.2, the window was extended by 50 bp; if not, the analysis proceeded to the next methylated CpG. Windows were continually extended until the proportion of methylated CpGs in the window fell below the 0.2 criteria. The location of methylation islands in the genome were characterized using BEDtools `intersect` v2.26.0.

### Differential Methylation Analysis

Differential methylation analysis for individual CpG dinucleotides was performed using `methylKit` v.1.7.9 in R (Akalin et al., 2012) using deduplicated, sorted bam files as input. Only CpGs with at least 5x coverage in each sample were considered for analysis. Methylation differences between treatments were obtained for all loci in the CpG background using `calculateDiffMeth`, a logistic regression built into `methylKit`. The logistic regression models the log odds ratio based on the proportion of methylation at each locus:

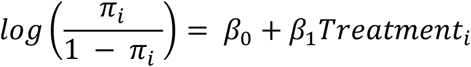

A differentially methylated locus (DML) was defined as an individual CpG dinucleotide with at least a 50% methylation change between treatment and control groups, and a q-value < 0.01 based on correction for false discovery rate with the SLIM method (Wang et al., 2011). Hypermethylated DML were defined as those with significantly higher percent methylation in oysters exposed to high *p*CO_2_ conditions, and hypomethylated DML with significantly lower percent methylation in the high *p*CO_2_ treatment. A Principal Components Analysis (PCA) was performed for differentially methylated loci (DML) for oyster sample methylation profiles between treatments, then compared to a PCA for all MBD-enriched CpG loci. The location of DML were characterized in relation to putative promoter regions, UTR, exons, introns, transposable elements, and intergenic regions using BEDtools `intersect` v2.26.0. Loci that did not overlap with the aforementioned genomic features were also identified. A chi-squared contingency test was used to test the null hypothesis of no association between genomic location and methylation status between MBD-enriched CpGs and DML. To describe the location of DML across different gene architectures, the position of DML in the gene was scaled from 0 to 100 bp.

### Enrichment Analysis

Functional enrichment analyses were used to determine if any biological processes were overrepresented in genes based on individual CpG methylation levels. Enrichment was conducted with GO-MWU, a rank-based gene enrichment method initially developed for analyzing transcriptomics data (Wright et al., 2015). Instead of only using genes with DML, GO-MWU identifies GO categories that are overrepresented by genes with any CpGs, allowing for more data to contribute to any trends. GO-MWU scripts and a gene ontology database were downloaded from the GO-MWU Github repository (https://github.com/z0on/GO_MWU).

A gene list and table of significance measures were used as GO-MWU analysis inputs. The gene list contained Genbank IDs and all associated gene ontology terms. For the table of significance measures, Genbank IDs were matched with the smallest *P*-value for associated CpGs analyzed by `methylKit`. To match the Genbank IDs to CpG loci within mRNAs and create the gene list, overlaps between the *C. virginica* mRNA track from NCBI and the CpG background used in `methylKit` were obtained using BEDtools `intersect` v2.26.0. The mRNAs were then annotated with Uniprot Accession codes using a BLASTx search (v.2.2.29; (Gish and States, 1993; UniProt Consortium, 2019). The Uniprot Swiss-Prot Database (downloaded from SwissProt 2018-06-15) was used to obtain protein information and Uniprot Accession codes. Genbank IDs provided by NCBI were used to match CpG background-mRNA overlaps with the annotated mRNA track. Gene ontology terms were paired to Uniprot Accession codes using the Uniprot Swiss-Prot Database (UniProt Consortium, 2019). All GO-MWU inputs are available in the associated Github repository (Venkataraman 2020).

Once analysis inputs were created, gene ontology terms for each gene were matched with parental terms using default GO-MWU settings. Parental ontology categories with the exact same gene list were combined. Groups were further combined if they shared at least 75% of the same genes. After clustering was complete, a Mann-Whitney U test identified gene ontology categories that were significantly enriched by corresponding hyper- or hypo-methylated loci in genes using the default 10% FDR. Genes with DML were mapped to gene ontology subsets (GO Slim terms) for biological processes to further categorize gene functions.

## Results

### Water Chemistry

All oysters were initially subjected to acclimation *p*CO_2_ conditions (*p*CO_2_ = 521 ± 32 ppm, Ω_calcite_= 2.82 ± 0.13) for 14 days. Following acclimation the treatments were initiated. Oysters in control *p*CO_2_ conditions (*p*CO_2_ = 492 ± 50 μatm; Ω_calcite_ = 3.01 ± 0.25) experienced low *p*CO_2_ and higher Ω_calcite_ than those in elevated *p*CO_2_ conditions (*p*CO_2_ = 2550 ± 211 μatm; Ω_calicite_ = 0.72 ± 0.06) (Table 1).

**Table 1.** Summary of water chemistry during the 14-day acclimation period and 28-day experimental exposure. Values indicate mean and standard error for temperature (T), salinity (S), dissolved inorganic carbon (DIC), total alkalinity (A_T_), calculated pH on seawater scale, calculated *p*CO_2_, and calculated calcite saturation (Ω_calcite_).

| Experimental Stage | T (°C) | S (PSU) | DIC | A<sub>T</sub> | pH<sub>sw</sub> | pCO<sub>2</sub> (μatm) | Ω<sub>calcite</sub> |
| --- | --- | --- | --- | --- | --- | --- | --- |
| Acclimation | 14.6 ± 0.4 | 31.4 ± 0.1 | 1978 ± 7 | 2127 ± 6 | 7.94 ± 0.00 | 521 ± 32 | 2.82 ± 0.13 |
| Control pCO<sub>2</sub> Conditions | 14.5 ± 0.4 | 31.6 ± 0.3 | 1960 ± 32 | 2140 ± 15 | 7.95 ± 0.01 | 492 ± 50 | 3.01 ± 0.25 |
| Elevated pCO<sub>2</sub> Conditions | 14.5 ± 0.3 | 31.54 ± 0.32 | 2173 ± 37 | 2132 ± 42 | 7.29 ± 0.01 | 2550 ± 211 | 0.72 ± 0.06 |

### MBD-BS-Seq

DNA sequencing yielded 280 million DNA sequence reads (NCBI Sequence Read Archive: BioProject accession number PRJNA513384). Of 276 million trimmed paired-end reads, 136 million (49.4%) were mapped to the *C. virginica* genome, providing an average of 13.6 million reads per sample. Sequencing efforts provided data for 4,304,257 CpG loci (30.7% of 14,458,703 total CpGs in the *C. virginica* genome) with at least 5x coverage across all samples combined. As expected, the location of CpGs with 5x coverage in the genome differed from the distribution of all CpG motifs (Contingency test; χ^2^ = 1,306,900, df = 6, *P*-value < 2.2e-16). Of all loci with 5x coverage, 3,255,049 CpGs (75.6%) were found in genic regions in 33,126 out of 38,929 annotated genes in the genome.

The general methylation landscape was defined using all loci with a minimum 5x coverage in each sample. The majority, 3,181,904 (73.9% of MBD-Enriched loci) loci were methylated, with 481,788 (11.2%) sparsely methylated loci and 640,565 (14.9%) unmethylated loci (Figure 1A). Median values for global percent methylation and sample methylation varied across genome features (Figure 1B-G). Based on these parameters and data, we calculated that 22% of all CpGs in the gonads (2.7% of total cytosines) had methylation levels greater than 50%. Loci methylation was characterized in relation to putative promoters, UTR, exons, introns, transposable elements, and intergenic regions (Figure 2). Methylated CpGs were found primarily in genic regions, with 2,521,653 loci (79.2%) in 25,496 genes. We rejected the null hypothesis that CpG methylation status was independent of genomic location, as the proportion of methylated CpG loci was different than expected in putative promoters, UTR, exons, introns, transposable elements, and intergenic regions (Contingency test; χ^2^ = 1,311,600, df = 6, *P*-value < 2.2e-16; Figure 2). There was a larger proportion of methylated loci found in exons compared to all CpGs in the genome (Figure 2). Methylated loci were also found in introns (with 1,448,786 loci (47.3% of methylated loci) versus 1,013,691 CpGs (31.9%) in exons), although this was not higher than expected based on the distribution of all CpGs. Transposable elements contained 755,222 methylated CpGs (23.7%). Putative promoter regions overlapped with 106,111 loci (3.3%), UTR with 128,585 loci (4.0%), and intergenic regions with 660,197 loci (20.7%). There were 372,047 methylated loci (11.7%) that did not overlap with either exons, introns, transposable elements, or promoter regions.

**Figure 1.**
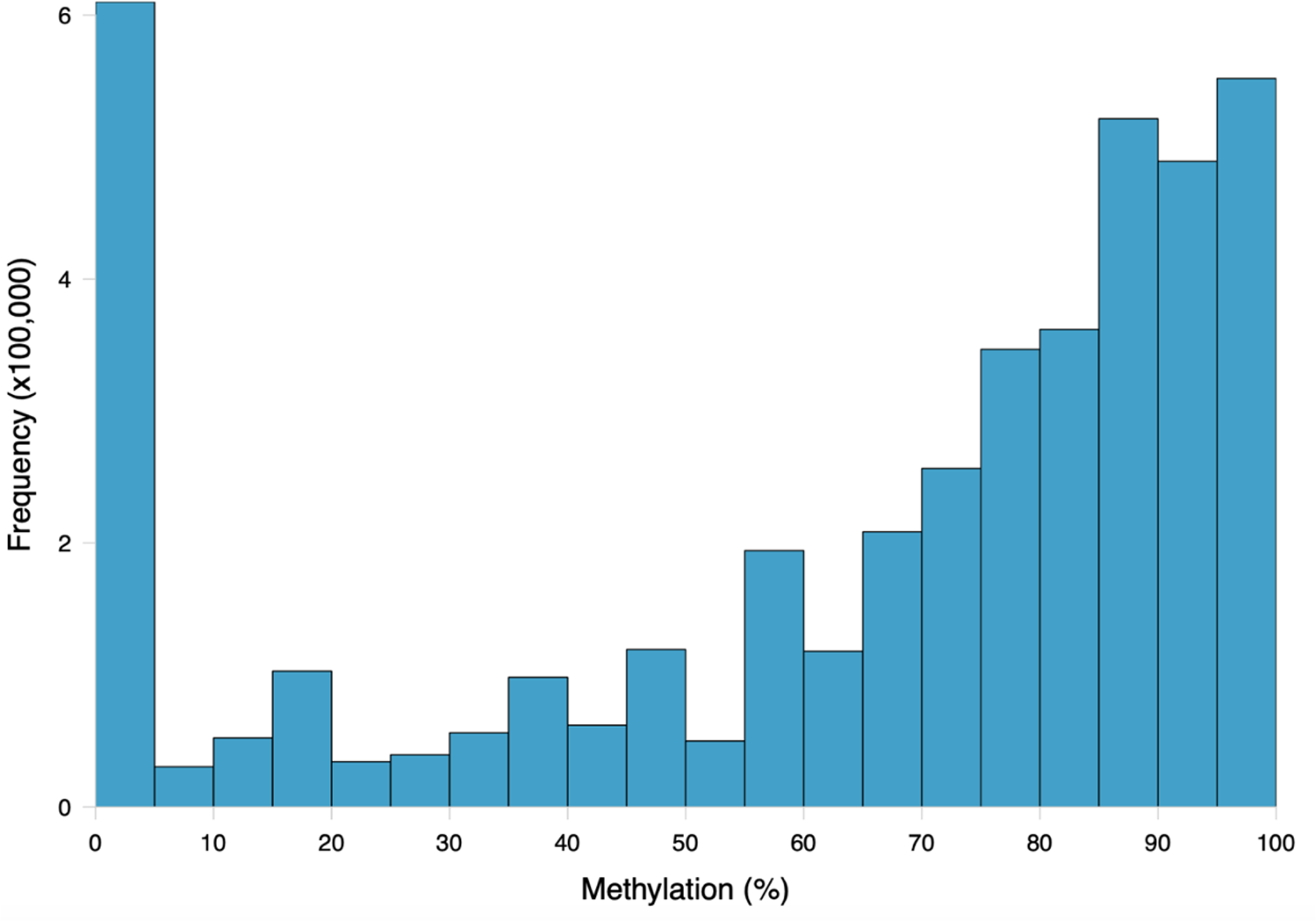
Frequency distribution of methylation ratios for CpG loci in *C. virginica* gonad tissue DNA subjected to MBD enrichment. A total of 4,304,257 CpGs with at least 5x coverage summed across all ten samples were characterized. Loci were considered methylated if they were at least 50% methylated, sparsely methylated loci were 10-50% methylated, and unmethylated loci were 0-10% methylated.

**Figure 2.**
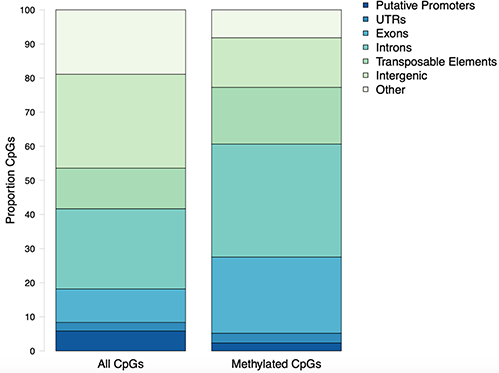
Proportion of CpG loci within genomic features. All CpGs are every dinucleotide in the *C. virginica* genome. Methylated CpGs refers to a dinucleotide with a methylation level of at least 50%.

A total of 37,063 methylation islands were identified in the *C. virginica* genome (Venkataraman et al., 2020). Methylation islands contained between 11 and 24,777 methylated CpGs, with a median of 30 methylated CpGs per island. Lengths of methylation islands ranged from 500 to 1,236,482 base pairs, with a median length of 1,024 base pairs. The majority of methylation islands (36,017; 97.2%) were less than 100,000 bp in length. There were 30,773 (83.0%) methylation islands that overlapped with genic regions.

### Differential Methylation Analysis

A total of 598 CpG loci were differentially methylated between oysters exposed to control or high *p*CO_2_, with 51.8% hypermethylated and 48.2% hypomethylated between treatments (Figure 3; Venkataraman 2020). When considering a PCA using methylation status of all CpG loci with 5x coverage across all samples, the first two principal components explained 29.8% of sample variation (Figure 4A). The first two principal components in a PCA with only differentially methylated loci (DML) explained 57.1% of the variation among treatments (Figure 4B). These DML were distributed throughout the *C. virginica* genome (Figure 5). The fifth chromosome had the most DML normalized by number of CpGs in the chromosome, and had the most genes; however, this was not the largest chromosome (Figure 5A).

**Figure 3.**
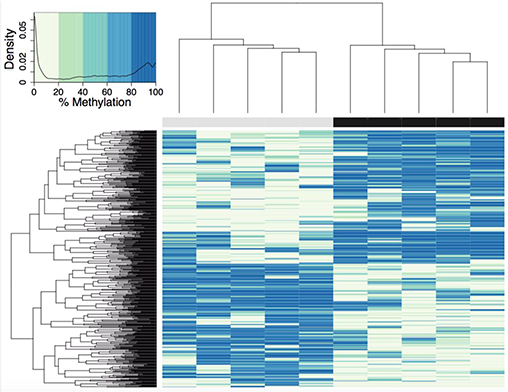
Heatmap of DML in *C. virginica* reproductive tissue created using a euclidean distance matrix. Samples in control *p*CO_2_ conditions are represented by grey, and samples in elevated *p*CO_2_ conditions are represented by a black bar. Loci with higher percent methylation are represented by darker colors. A logistic regression identified 598 DML, defined as individual CpG dinucleotide with at least a 50% methylation change between treatment and control groups, and a q-value < 0.01 based on correction for false discovery rate with the SLIM method. The density of DML at each percent methylation value is represented in the heatmap legend.

**Figure 4.**
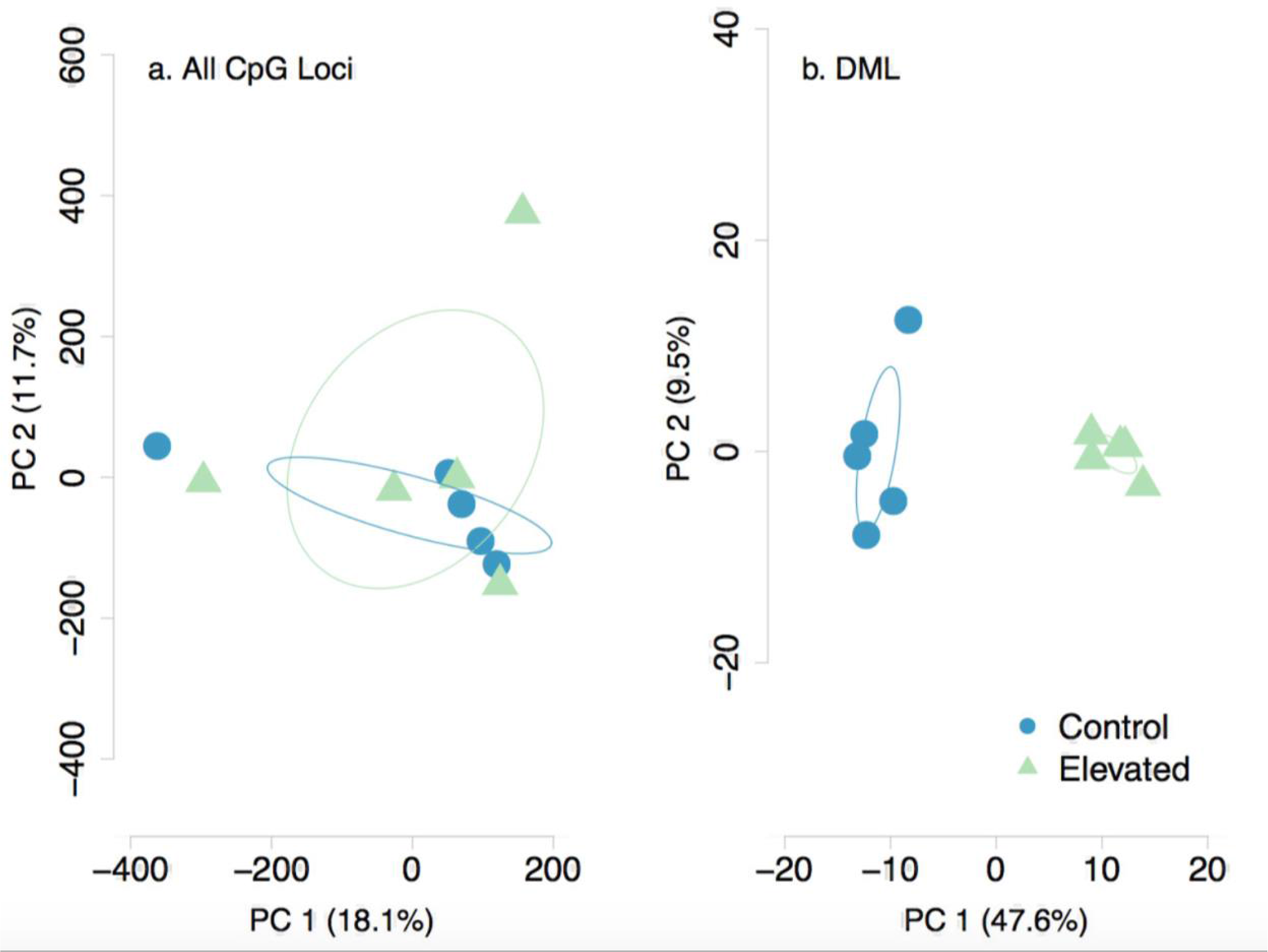
Principal Components Analysis of a) all CpG loci with 5x coverage across samples and b) DML. Methylation status of individual CpG loci explained 29.2% of variation between samples when considering all CpG loci. Methylation status of DML explained 57.1% of sample variation.

**Figure 5.**
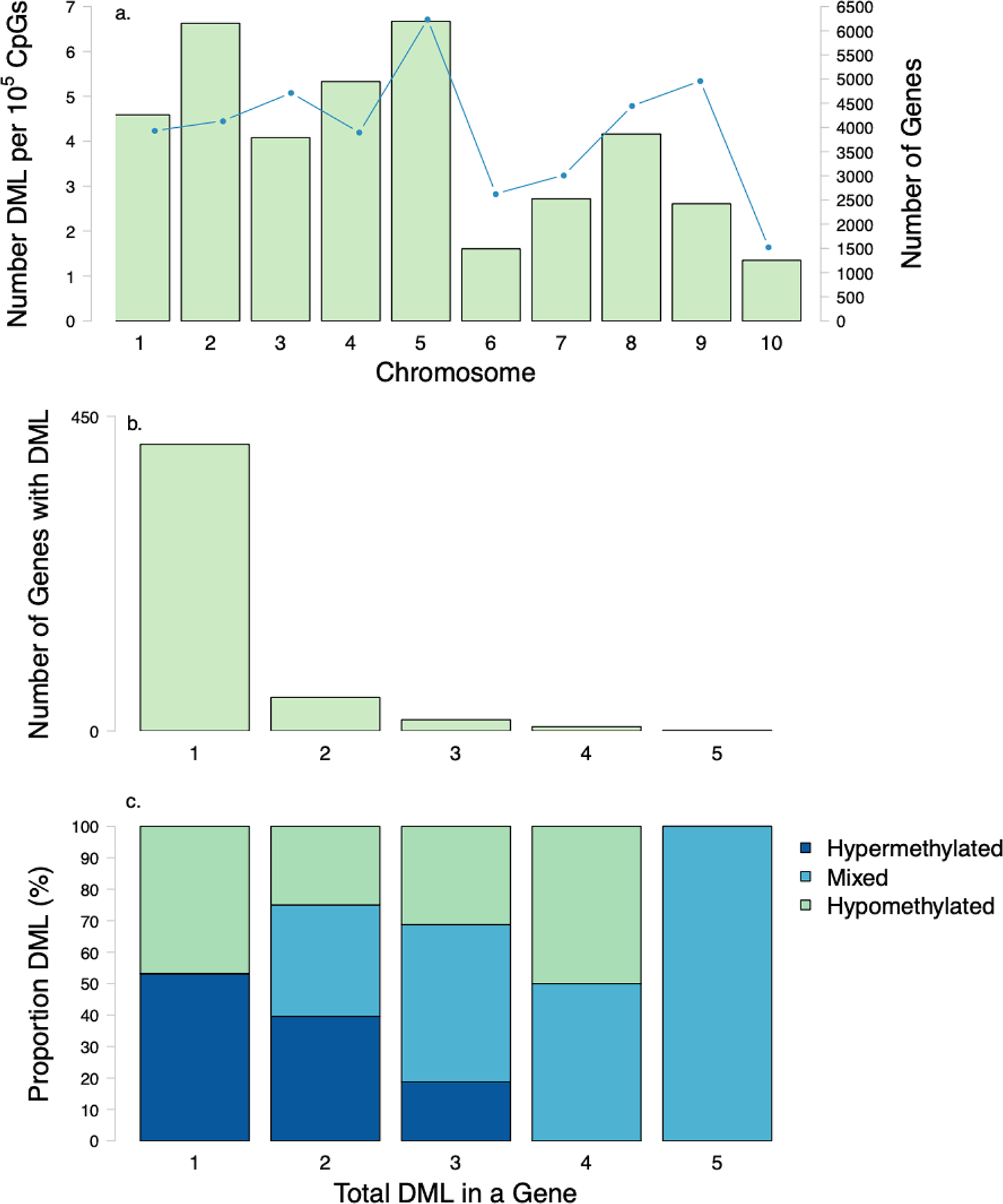
Distribution of DML among chromosomes and genes. (a) Number of DML normalized by number of CpG in each chromosome (bars) and number of genes (line) in each *C. virginica* chromosome. (b) Number of genes with various numbers of DML per gene (1-5). Most genes that contained DML only had 1 DML. (c) Proportion of hypermethylated, hypomethylated DML in genes with various numbers of DML per gene (1-5). Mixed refers to a classification of a gene that has both hypermethylated and hypomethylated DML.

Examination of DML within genes revealed that some genes contained multiple DML (Figure 5B-C). Of the 481 genes with DML, the majority only contained one DML (Figure 5B). There were 48 genes with 2 DML, 16 genes with 3 DML, 6 genes with 4 DML and 1 gene with 5 DML (Figure 5B). When multiple DML were found within a gene, they could be methylated in either the same or opposite directions (Figure 5C).

Within the genome, DML were mostly present in genic regions, with 560 DML in 481 genes (368 DML in exons and 192 in introns). In addition, 42 DML were found in putative promoter regions, 27 in UTR, 57 in transposable elements, and 38 in intergenic regions. There were 21 DML located outside of exons, introns, transposable elements, and putative promoters. Additionally, 537 DML were found in methylation islands. The distribution of DML in *C. virginica* gonad tissue was higher in exons than expected for MBD-enriched CpG loci with minimum 5x coverage across all samples (Contingency test; χ^2^ = 401.09, df = 6, *P*-value < 2.2e-16; Figure 6). Of the 598 DML, 310 were hypermethylated and 288 were hypomethylated in the high *p*CO2 treatment. The number of hyper- and hypomethylated DML was almost evenly split within each genomic feature, with the exception of putative promoter regions that had 44 hypermethylated DML versus 23 hypomethylated DML. Within a gene, DML did not appear to be concentrated in one particular region. The distribution of hyper- and hypomethylated DML along a gene do not differ from each other (Figure 7).

**Figure 6.**
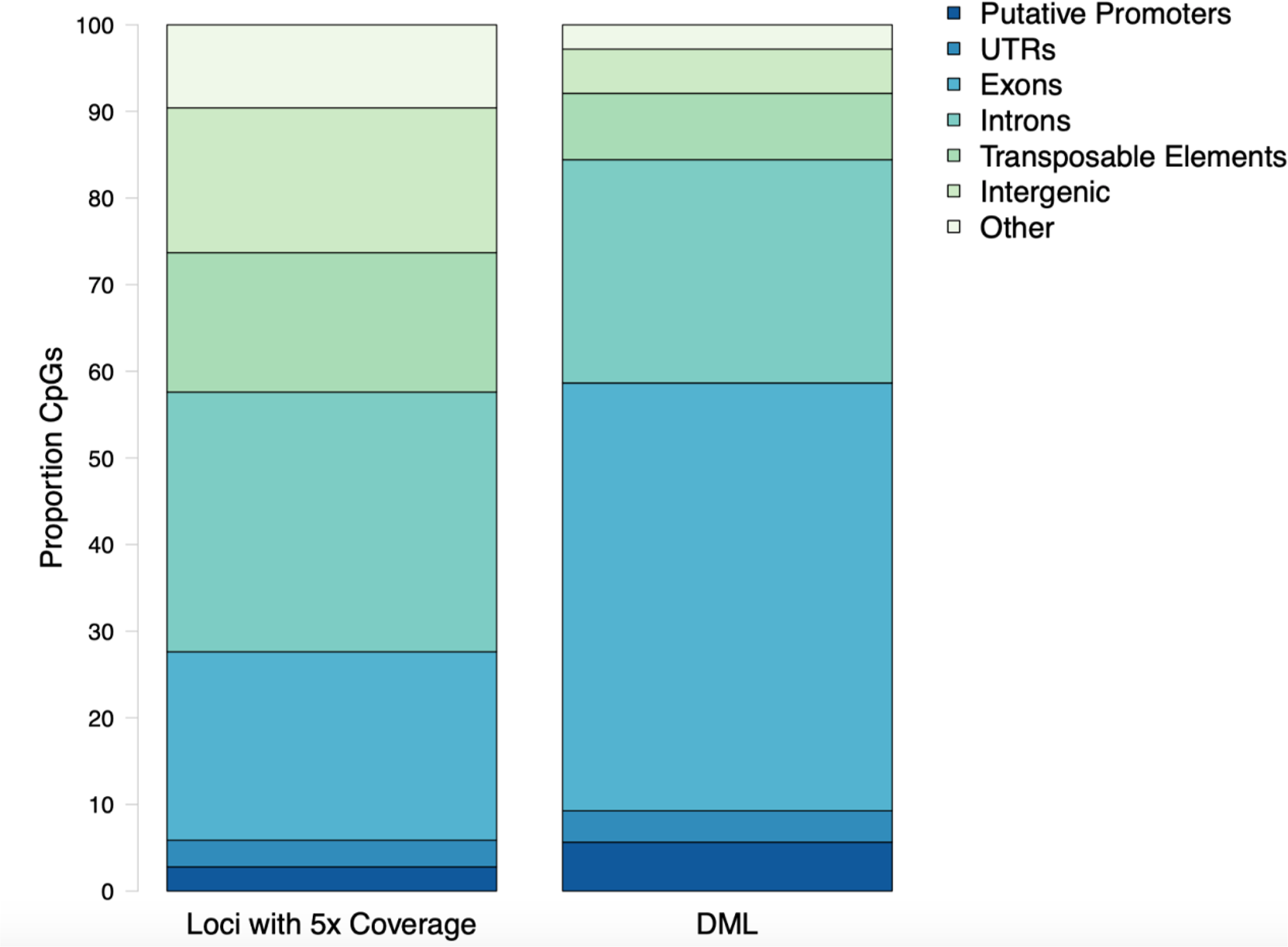
Proportion CpG loci within putative promoters, untranslated regions (UTR), exons, introns, transposable elements, and intergenic regions for MBD-enriched CpGs and differentially methylated loci (DML). The distribution of DML in *C. virginica* gonad tissue in response to ocean acidification differed from distribution of MBD-enriched loci with 5x coverage across control and treatment samples (Contingency test; χ^2^ = 401.09, df = 6, *P*-value < 2.2e-16).

**Figure 7.**
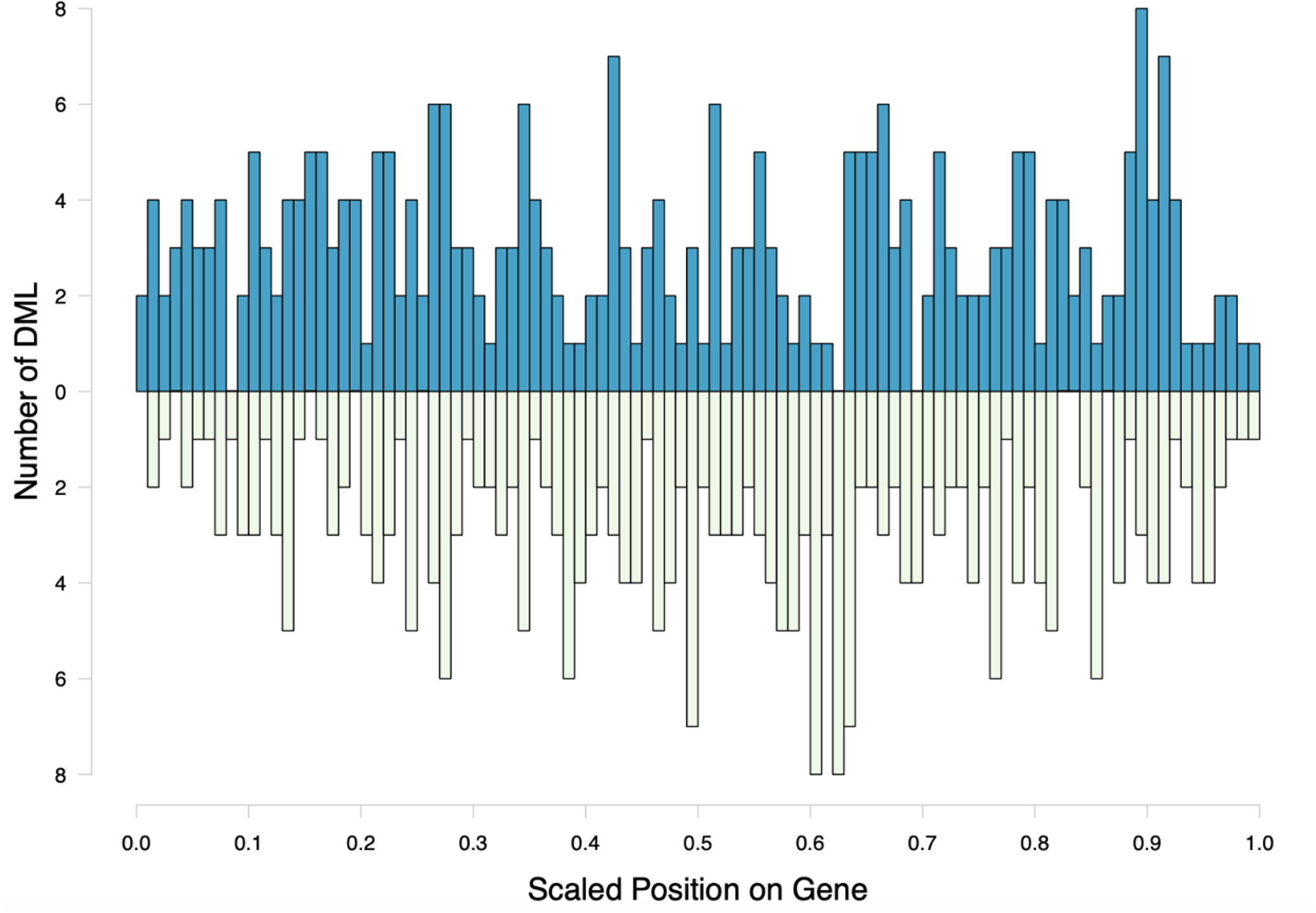
Distribution of hyper- and hypomethylated DML along a hypothetical gene. The scaled position of a DML within a gene was calculated by dividing the base pair position of the DML by gene length. Counts of hypermethylated DML are plotted above the x-axis, and hypomethylated DML counts are below the x-axis.

The DML were found in genes responsible for various biological processes. However, no gene ontology categories were significantly represented (Figure 8). The majority of genes with DML were involved in protein ubiquitination processes. These genes were not consistently hyper- or hypomethylated. Certain biomineralization genes did contain DML. The gene coding for calmodulin-regulated spectrin-associated protein contained three hypomethylated and one hypermethylated DML. Genes coding for EF-hand protein with calcium-binding domain, calmodulin-binding transcription activator, and calmodulin-lysine N-methyltransferase contained one or two hypermethylated DML.

**Figure 8.**
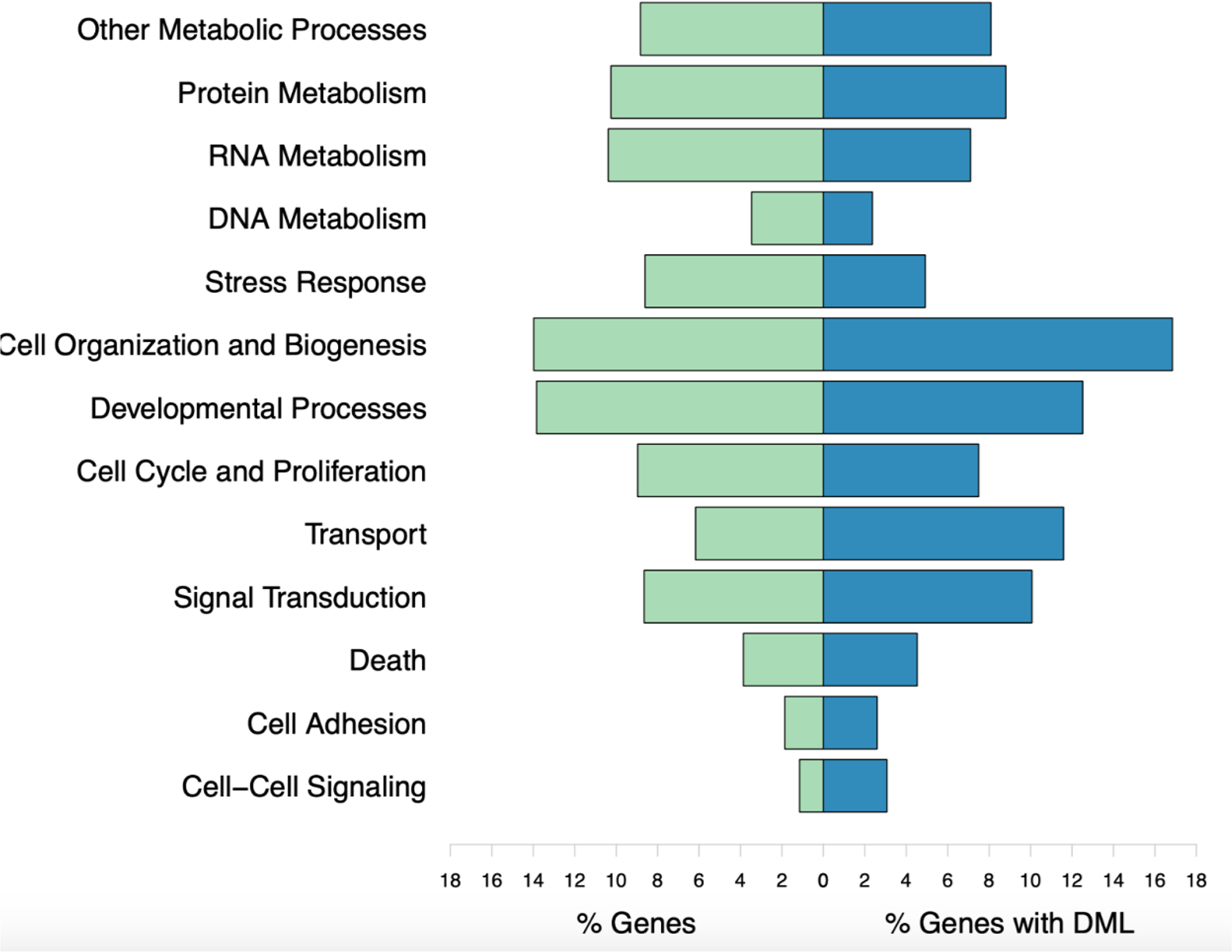
Biological processes represented by all genes used in enrichment background (% Genes) and those with DML (% Genes with DML). Gene ontology categories with similar functions are represented by the same color. Genes may be involved in multiple biological processes. No gene ontologies were significantly enriched.

## Discussion

The present study is a general description of DNA methylation in *C. virginica*, and is one of the first to examine epigenetic responses to ocean acidification in the gonad tissue of a mollusc species. Five hundred ninety-eight differentially methylated loci (DML) were identified in response to the elevated *p*CO_2_ treatments, most of which were in exons. Not only was DNA methylation of *C. virginica* altered in response to ocean acidification, but changes in gonad methylation indicates potential for these methylation patterns to be inherited by offspring.

Understanding how environmental stressors influence the epigenome is crucial when considering potential acclimatization mechanisms in marine invertebrates. Our finding that high *p*CO_2_ impacts *C. virginica* DNA methylation adds to a growing body of work about ocean acidification’s impact on marine invertebrate methylomes. The coral species *P. damicornis* demonstrated an overall increase in DNA methylation when exposed to low pH conditions (7.3 - 7.6) for six weeks, potentially influencing biomineralization (Putnam et al., 2016). Another coral species, *S. pistillata*, also demonstrated an increase in genome-wide DNA methylation when exposed to low pH conditions for two years. Changes in the methylome also modified gene expression and altered pathways involved in cell cycle regulation (Liew et al., 2018b). The present study on an oyster, however, did not observe the overall genome-wide increase in methylation that was reported for corals. Instead, we found subtle, but significant, increases or decreases in percent methylation at several hundred individual CpGs distributed across the genome. As *C. virginica* and coral species are adapted to different environments and ecological niches, it is possible that species-specific differences in methylation responses contribute to the observed methylation pattern.

The *C. virginica* methylation landscape suggests a role for methylation in gene activity. Approximately 22% of CpGs in the *C. virginica* gonad genome were methylated, which is consistent with previous studies of marine invertebrate genomes Gavery and Roberts, 2013; Olson and Roberts, 2014; Hofmann, 2017; Dimond and Roberts, 2020). Methylated loci were concentrated in introns for *C. virginica*, followed by exons and transposable elements. This location of methylated CpGs in gene bodies is consistent with what has been reported across similar taxa (Roberts and Gavery, 2012; Eirin-Lopez and Putnam, 2018). The concentration of methylated CpGs in gene bodies corresponds with proposed functionality in influencing gene activity (Roberts and Gavery, 2012; Dixon et al., 2014; Liew et al., 2018b). Our study also found methylation in transposable elements, putative promoters and intragenic regions. In plants, transposable element methylation has been shown to modulate the effect of transposable element insertion in genic regions (Hosaka and Kakutani, 2018). It is possible that methylation of transposable elements in *C. virginica* could also limit the effect of transposable elements. The characterization of methylation islands in the *C. virginica* genome demonstrates the viability of this descriptive tool for future work examining methylation in mollusc species.

The presence of DML suggests that exposure to experimental ocean acidification conditions elicits an epigenetic response. Many studies have documented changes to oyster protein synthesis, energy production, metabolism, antioxidant responses, and reproduction in response to ocean acidification (Tomanek et al., 2011; Timmins-Schiffman et al., 2014; Dineshram et al., 2016; Boulais et al., 2017; Omoregie et al., 2019). Examination of methylation associated with these physiological responses could identify mechanisms that contribute to these changes. For example, our study found a hypomethylated DML in the heat shock protein 75 kDA gene, and gene expression responses to ocean acidification in *C. virginica* have found downregulation in a similar molecular chaperone, heat shock protein 70kDa (Beniash et al., 2010; Ivanina et al., 2014). Other gene expression studies in bivalves have found changes in oxidative stress proteins such as superoxide dismutase, cytochrome c, peroxiredoxin, and NADH dehydrogenase (Chapman et al., 2011; Clark et al., 2013; Goncalves et al., 2016, 2017). Although we did not find any DML in these genes, combined study of DNA methylation and transcription may reveal how changes in gene expression are regulated in response to environmental stressors.

Although DML were found across various genome features, they were mostly in exons and introns. This is consistent with a recent study of *C. virginica* gill tissue found differentially methylated regions in response to a salinity gradient were primarily in genic regions ((Johnson and Kelly, 2019). Interestingly, DML were not found consistently in one particular region of a gene. Similarly, methylated positions in genic regions were evenly distributed after the coral *S. pistillata* was exposed to low pH (Liew et al., 2018b). Examination of another coral, *P. daedalea*, in different temperature and salinity conditions found more frequent methylation at 5’ and 3’ ends of genes (Liew et al., 2018a). We also found several genes with multiple DML. These DML were not consistently hyper- or hypomethylated in the same gene. As hyper- and hypomethylation may result in different transcriptional outcomes, future work should examine the role of multiple DML on alternative splicing and gene expression.

The concentration of DML in gene bodies suggests a role for DNA methylation in gene expression and regulation. A majority of genes with DML were involved in protein ubiquitination. Protein ubiquitination is a post-translational protein modification that is involved in protein synthesis and degradation (Peng et al., 2003; Komander, 2009). Previous studies in which oysters were exposed to experimental ocean acidification conditions have demonstrated changes in this pathway. For example, shotgun proteomic characterization of posterior gill lamellae from adult *C. gigas* exposed to high *p*CO_2_ revealed increased abundance of proteins involved in ubiquitination and decreased protein degradation (Timmins-Schiffman et al., 2014). Elevated *p*CO_2_ levels were also found to upregulate malate dehydrogenase in adult *C. virginica* mantle tissue (Tomanek et al., 2011). Several genes involved in protein ubiquitination, including those for malate dehydrogenase, ubiquitin-protein ligase, RNA polymerase-associated protein, and DNA damage-binding protein, were significantly hypermethylated in gonad tissue exposed to elevated *p*CO_2_. Hypermethylation of these genes may decrease transcriptional opportunities, thus indicating a critical role in the response to ocean acidification.

Four genes involved in biomineralization contained DML, suggesting these genes can be epigenetically regulated. Upregulation of calcium-binding gene expression has been previously documented in *C. virginica* (Richards et al., 2018). Since the hypermethylated DML in these genes are typically associated with reduced transcriptional opportunities, it is unclear how methylation changes relate to gene expression for biomineralization genes. Many studies examining ocean acidification-induced carryover effects in bivalves note changes to calcification processes. For example, the Sydney rock oyster (*S, glomerata*) larvae exhibit faster shell growth in high *p*CO_2_ conditions when parents mature in those same conditions (Parker et al., 2012, 2015). In contrast, larvae from other species found in the North Atlantic such as northern quahog (hard clam; *M. mercenaria*) and bay scallops (*A. irradians*) developed slower when parents were reproductively conditioned in low pH conditions (Griffith and Gobler, 2017). There is some evidence to suggest that *C. virginica* larvae may be more resilient to high *p*CO_2_ conditions than *M. mercenaria* or *A. irradians (Gobler and Talmage, 2014)*. Differential methylation of biomineralization genes in *C. virginica* reproductive tissue could be a mechanism to explain when parental experience impacts larval calcification if in fact these DML are inherited.

Although our work documents significant changes to DNA methylation in reproductive tissue after high *p*CO_2_ exposure, this finding may be confounded by secondary effects of gonad maturation. Specimens collected were from mixed populations, and sampled tissue contained both mature and immature gametes. Reproductive tissue likely contained both gametic and somatic cell types. Sex-specific effects have also been documented in response to ocean acidification in mollusc species (Parker et al., 2018; Venkataraman et al., 2019). Lack of a reproductive phenotype precludes any interpretation of how maturation stage or sex can influence changes DNA methylation, as previous work in *C. gigas* demonstrates these factors as significant influences on baseline methylation patterns (Zhang et al., 2018). Nevertheless, differential methylation in stress response and biomineralization genes suggests that our study does record epigenetic responses to ocean acidification. Future work should pair methylation data with reproductive phenotypes to provide additional information on sex- or stage-specific epigenetic responses to ocean acidification.

## Conclusion

Our study found that *C. virginica* demonstrates a significant epigenetic response to elevated *p*CO_2_ exposure, with 598 DML identified. The concentration of these DML in gene bodies suggests that methylation may be important for transcriptional control in response to environmental stressors. As ocean acidification induced differential methylation in *C. virginica* gonad tissue, there is a potential for intergenerational epigenetic inheritance, which could control the gene activity of processes such as biomineralization. As carryover effects can persist even when stressors are long-removed ((Venkataraman et al., 2019)), understanding the mechanisms involved in intergenerational acclimatization is crucial. Future work should focus on methylation patterns in adult *C. virginica* fully-formed gametes and larvae exposed to various *p*CO_2_ conditions to determine to what degree a difference in methylation influences gene activity and how this might influence phenotypic plasticity.

## Acknowledgements

This project was funded by National Science Foundation Biological Oceanography award 1635423 to KEL, JR, and SBR, and a Hall Conservation Genetics Research Award to YRV. This work was facilitated through the use of advanced computational, storage, and networking infrastructure provided by the Hyak supercomputer system at the University of Washington. We also thank Mackenzie Gavery and our two reviewers for their insightful feedback on the manuscript.

## Data Accessibility

Raw sequence data is available at the NCBI Sequence Read Archive under BioProject accession number PRJNA513384, with associated metadata and information also available at Woods Hole Open Access Server: https://hdl.handle.net/1912/25138

Associated information for all analyses and supplemental material can be found in the Github repository which is available in an archival format (Venkataraman 2020; https://doi.org/10.6084/m9.figshare.11923479)

## References

Akalin, A., Kormaksson, M., Li, S., Garrett-Bakelman, F. E., Figueroa, M. E., Melnick, A., et al. (2012). methylKit: a comprehensive R package for the analysis of genome-wide DNA methylation profiles. Genome Biol. 13, R87.

Andrews, S. (2010). FastQC: a quality control tool for high throughput sequence data.

Bao, W., Kojima, K. K., and Kohany, O. (2015). Repbase Update, a database of repetitive elements in eukaryotic genomes. Mob. DNA 6, 11.

Barton, A., Hales, B., Waldbusser, G. G., Langdon, C., and Feely, R. A. (2012). The Pacific oyster, *Crassostrea gigas*, shows negative correlation to naturally elevated carbon dioxide levels: Implications for near-term ocean acidification effects. Limnol. Oceanogr. 57, 698–710.

Beniash, E., Ivanina, A., Lieb, N. S., Kurochkin, I., and Sokolova, I. M. (2010). Elevated level of carbon dioxide affects metabolism and shell formation in oysters Crassostrea virginica. Mar. Ecol. Prog. Ser. 419, 95–108.

Bird, A. (2002). DNA methylation patterns and epigenetic memory. Genes Dev. 16, 6–21.

Bossdorf, O., Richards, C. L., and Pigliucci, M. (2008). Epigenetics for ecologists. Ecol. Lett. 11, 106–115.

Boulais, M., Chenevert, K. J., Demey, A. T., Darrow, E. S., Robison, M. R., Roberts, J. P., et al. (2017). Oyster reproduction is compromised by acidification experienced seasonally in coastal regions. Sci. Rep. 7, 13276.

Byrne, M., Foo, S. A., Ross, P. M., and Putnam, H. M. (2019). Limitations of cross and multigenerational plasticity for marine invertebrates faced with global climate change. Glob. Chang. Biol. doi:10.1111/gcb.14882.

Chapman, R. W., Mancia, A., Beal, M., Veloso, A., Rathburn, C., Blair, A., et al. (2011). The transcriptomic responses of the eastern oyster, *Crassostrea virginica*, to environmental conditions. Mol. Ecol. 20, 1431–1449.

Clark, M. S., Thorne, M. A. S., Amaral, A., Vieira, F., Batista, F. M., Reis, J., et al. (2013). Identification of molecular and physiological responses to chronic environmental challenge in an invasive species: the Pacific oyster, *Crassostrea gigas*. Ecol. Evol. 3, 3283–3297.

Deans, C., and Maggert, K. A. (2015). What Do You Mean, “Epigenetic”? Genetics 199, 887–896.

Dickson, A. G. (1990). Standard potential of the reaction : AgCl_(s)_ + 12H2_(g)_ = Ag_(s)_ + HCl_(aq)_, and and the standard acidity constant of the ion HSO_4−_ in synthetic sea water from 273.15 to 318.15 K. The Journal of Chemical Thermodynamics 22, 113–127.

Dimond, J. L., and Roberts, S. B. (2016). Germline DNA methylation in reef corals: patterns and potential roles in response to environmental change. Mol. Ecol. 25, 1895–1904.

Dimond, J. L., and Roberts, S. B. (2020). Convergence of DNA Methylation Profiles of the Reef Coral *Porites astreoides* in a Novel Environment. Frontiers in Marine Science 6, 792.

Dineshram, R., Chandramouli, K., Ko, G. W. K., Zhang, H., Qian, P.-Y., Ravasi, T., et al. (2016). Quantitative analysis of oyster larval proteome provides new insights into the effects of multiple climate change stressors. Glob. Chang. Biol. 22, 2054–2068.

Dixon, G. B., Bay, L. K., and Matz, M. V. (2014). Bimodal signatures of germline methylation are linked with gene expression plasticity in the coral *Acropora millepora*. BMC Genomics 15, 1109.

Eirin-Lopez, J. M., and Putnam, H. M. (2018). Marine Environmental Epigenetics. Ann. Rev. Mar. Sci. doi:10.1146/annurev-marine-010318-095114.

Ekstrom, J. A., Suatoni, L., Cooley, S. R., Pendleton, L. H., Waldbusser, G. G., Cinner, J. E., et al. (2015). Vulnerability and adaptation of US shellfisheries to ocean acidification. Nat. Clim. Chang. 5, 207–214.

Feely, R. A., Alin, S. R., Newton, J., Sabine, C. L., Warner, M., Devol, A., et al. (2010). The combined effects of ocean acidification, mixing, and respiration on pH and carbonate saturation in an urbanized estuary. Estuarine, Coastal and Shelf Science 88, 442–449. doi:10.1016/j.ecss.2010.05.004.

Gatzmann, F., Falckenhayn, C., Gutekunst, J., Hanna, K., Raddatz, G., Carneiro, V. C., et al. (2018). The methylome of the marbled crayfish links gene body methylation to stable expression of poorly accessible genes. Epigenetics Chromatin 11, 57.

Gavery, M. R., and Roberts, S. B. (2013). Predominant intragenic methylation is associated with gene expression characteristics in a bivalve mollusc. PeerJ 1, e215.

Gazeau, F., Quiblier, C., Jansen, J. M., Gattuso, J.-P., Middelburg, J. J., and Heip, C. H. R. (2007). Impact of elevated CO_2_ on shellfish calcification. Geophys. Res. Lett. 34, L07603.

Gish, W., and States, D. J. (1993). Identification of protein coding regions by database similarity search. Nat. Genet. 3, 266–272.

Gobler, C. J., and Talmage, S. C. (2014). Physiological response and resilience of early life-stage Eastern oysters (*Crassostrea virginica*) to past, present and future ocean acidification. Conserv Physiol 2, cou004.

Gómez-Chiarri, M., Warren, W. C., Guo, X., and Proestou, D. (2015). Developing tools for the study of molluscan immunity: The sequencing of the genome of the eastern oyster, *Crassostrea virginica*. Fish & Shellfish Immunology 46, 2–4. doi:10.1016/j.fsi.2015.05.004.

Goncalves, P., Anderson, K., Thompson, E. L., Melwani, A., Parker, L. M., Ross, P. M., et al. (2016). Rapid transcriptional acclimation following transgenerational exposure of oysters to ocean acidification. Mol. Ecol. 25, 4836–4849.

Goncalves, P., Thompson, E. L., and Raftos, D. A. (2017). Contrasting impacts of ocean acidification and warming on the molecular responses of CO_2_-resilient oysters. BMC Genomics 18, 431.

Griffith, A. W., and Gobler, C. J. (2017). Transgenerational exposure of North Atlantic bivalves to ocean acidification renders offspring more vulnerable to low pH and additional stressors. Sci. Rep. 7, 11394.

Helm, M. M., and Bourne, N. (2004). Hatchery Culture of Bivalves: A Practical Manual. Food & Agriculture Org.

Hofmann, G. E. (2017). Ecological Epigenetics in Marine Metazoans. Frontiers in Marine Science 4. doi:10.3389/fmars.2017.00004.

Hosaka, A., and Kakutani, T. (2018). Transposable elements, genome evolution and transgenerational epigenetic variation. Curr. Opin. Genet. Dev. 49, 43–48.

IPCC, 2019: Summary for Policymakers. In: IPCC Special Report on the Ocean and Cryosphere in a Changing Climate [H.-O. Pörtner, D.C. Roberts, V. Masson-Delmotte, P. Zhai, M. Tignor, E. Poloczanska, K. Mintenbeck, M. Nicolai, A. Okem, J. Petzold, B. Rama, N. Weyer (eds.)]. In press.

Ivanina, A. V., Hawkins, C., and Sokolova, I. M. (2014). Immunomodulation by the interactive effects of cadmium and hypercapnia in marine bivalves *Crassostrea virginica* and *Mercenaria mercenaria*. Fish Shellfish Immunol. 37, 299–312.

Jeong, H., Wu, X., Smith, B., and Yi, S. V. (2018). Genomic Landscape of Methylation Islands in Hymenopteran Insects. Genome Biol. Evol. 10, 2766–2776.

Johnson, K. M., and Kelly, M. W. (2019). Population epigenetic divergence exceeds genetic divergence in the Eastern oyster *Crassostrea virginica* in the Northern Gulf of Mexico. Evol. Appl. doi:10.1111/eva.12912.

Komander, D. (2009). The emerging complexity of protein ubiquitination. Biochem. Soc. Trans. 37, 937–953.

Krueger, F., and Andrews, S. R. (2011). Bismark: a flexible aligner and methylation caller for Bisulfite-Seq applications. Bioinformatics 27, 1571–1572.

Kurihara, H., Kato, S., and Ishimatsu, A. (2007). Effects of increased seawater *p*CO_2_ on early development of the oyster *Crassostrea gigas*. Aquat. Biol. 1, 91–98.

Langmead, B., and Salzberg, S. L. (2012). Fast gapped-read alignment with Bowtie 2. Nat. Methods 9, 357–359.

Lee, Kitach Tae-Wook, Kim Byrne, Robert H Millero, Frank J Feely, Richard A Liu, Yong-Ming (2010). The universal ratio of boron to chlorinity for the North Pacific and North Atlantic oceans. Geochimica et Cosmochimica Acta 74, 1801–1811.

Lewis, E., and Wallace, D. W. (1998). R: Program developed for CO_2_ system calculations ORNL/CDIAC-105. Carbon Dioxide Information Analysis CentreOak Ridge National Laboratory, US Department of Energy, Oak Ridge, Tennessee.

Liew, Y. J., Howells, E. J., Wang, X., Michell, C. T., Burt, J. A., Idaghdour, Y., et al. (2018a). Intergenerational epigenetic inheritance in reef-building corals. bioRxiv, 269076. doi:10.1101/269076.

Liew, Y. J., Zoccola, D., Li, Y., Tambutté, E., Venn, A. A., and Michell, C. T. (2018b). Epigenome-associated phenotypic acclimatization to ocean acidification in a reef-building coral. Science Advances.

Li, H., Handsaker, B., Wysoker, A., Fennell, T., Ruan, J., Homer, N., et al. (2009). The Sequence Alignment/Map format and SAMtools. Bioinformatics 25, 2078–2079.

Martin, M. (2011). Cutadapt removes adapter sequences from high-throughput sequencing reads. EMBnet.journal 17, 10–12.

Olson, C. E., and Roberts, S. B. (2014). Genome-wide profiling of DNA methylation and gene expression in Crassostrea gigas male gametes. Front. Physiol. 5, 224.

Omoregie, E., Mwatilifange, N. S. I., and Liswaniso, G. (2019). Futuristic Ocean Acidification Levels Reduce Growth and Reproductive Viability in the Pacific Oyster (*Crassostrea gigas*). J. Appl. Sci. Environ. Manage. 23, 1747–1754.

Parker, L. M., O’Connor, W. A., Byrne, M., Coleman, R. A., Virtue, P., Dove, M., et al. (2017). Adult exposure to ocean acidification is maladaptive for larvae of the Sydney rock oyster *Saccostrea glomerata* in the presence of multiple stressors. Biol. Lett. 13, 20160798.

Parker, L. M., O’Connor, W. A., Byrne, M., Dove, M., Coleman, R. A., Pörtner, H.-O., et al. (2018). Ocean acidification but not warming alters sex determination in the Sydney rock oyster, *Saccostrea glomerata*. Proc. R. Soc. B 285, 20172869.

Parker, L. M., O’Connor, W. A., Raftos, D. A., Pörtner, H.-O., and Ross, P. M. (2015). Persistence of Positive Carryover Effects in the Oyster, *Saccostrea glomerata*, following Transgenerational Exposure to Ocean Acidification. PLoS One 10, e0132276.

Parker, L. M., Ross, P. M., O’Connor, W. A., Borysko, L., Raftos, D. A., and Pörtner, H.-O. (2012). Adult exposure influences offspring response to ocean acidification in oysters. Glob. Chang. Biol. 18, 82–92.

Parker, L. M., Ross, P. M., O’Connor, W. A., Pörtner, H. O., Scanes, E., and Wright, J. M. (2013). Predicting the response of molluscs to the impact of ocean acidification. Biology 2, 651–692.

Peng, J., Schwartz, D., Elias, J. E., Thoreen, C. C., Cheng, D., Marsischky, G., et al. (2003). A proteomics approach to understanding protein ubiquitination. Nat. Biotechnol. 21, 921–926.

Putnam, H. M., Davidson, J. M., and Gates, R. D. (2016). Ocean acidification influences host DNA methylation and phenotypic plasticity in environmentally susceptible corals. Evol. Appl. 9, 1165–1178.

Quinlan, A. R., and Hall, I. M. (2010). BEDTools: a flexible suite of utilities for comparing genomic features. Bioinformatics 26, 841–842.

Richards, M., Xu, W., Mallozzi, A., Errera, R. M., and Supan, J. (2018). Production of Calcium-Binding Proteins in *Crassostrea virginica* in Response to Increased Environmental CO_2_ Concentration. Frontiers in Marine Science 5, 203.

Ries, J. B. (2011). A physicochemical framework for interpreting the biological calcification response to CO_2_-induced ocean acidification. Geochim. Cosmochim. Acta 75, 4053–4064.

Roberts, S. B., and Gavery, M. R. (2012). Is There a Relationship between DNA Methylation and Phenotypic Plasticity in Invertebrates? Front. Physiol. 2. doi:10.3389/fphys.2011.00116.

Rondon, R., Grunau, C., Fallet, M., Charlemagne, N., Sussarellu, R., Chaparro, C., et al. (2017). Effects of a parental exposure to diuron on Pacific oyster spat methylome. Environ Epigenet 3. doi:10.1093/eep/dvx004.

Ross, P. M., Parker, L., and Byrne, M. (2016). Transgenerational responses of molluscs and echinoderms to changing ocean conditions. ICES Journal of Marine Science. doi:10.1093/icesjms/fsv254.

Roy, Rabindra N Roy, Lakshimi N Vogel, Kathleen M Porter-Moore, C Pearson, Tara Good, Catherine E Millero, Frank J Campbell, Douglas M (1993). The dissociation constants of carbonic acid in seawater at salinities 5 to 45 and temperatures 0 to 45°C. Marine Chemistry 44, 249–267.

Smit, A. F. A., Hubley, R., and Green, P. (2013). 2013--2015. RepeatMasker Open-4.0.

Strader, M. E., Wong, J. M., Kozal, L. C., Leach, T. S., and Hofmann, G. E. (2019). Parental environments alter DNA methylation in offspring of the purple sea urchin, *Strongylocentrotus purpuratus*. J. Exp. Mar. Bio. Ecol. 517, 54–64.

Suzuki, M. M., and Bird, A. (2008). DNA methylation landscapes: provocative insights from epigenomics. Nat. Rev. Genet. 9, 465–476.

Timmins-Schiffman, E., Coffey, W. D., Hua, W., Nunn, B. L., Dickinson, G. H., and Roberts, S. B. (2014). Shotgun proteomics reveals physiological response to ocean acidification in *Crassostrea gigas*. BMC Genomics 15, 951.

Tomanek, L., Zuzow, M. J., Ivanina, A. V., Beniash, E., and Sokolova, I. M. (2011). Proteomic response to elevated PCO2 level in eastern oysters, *Crassostrea virginica*: evidence for oxidative stress. J. Exp. Biol. 214, 1836–1844.

UniProt Consortium (2019). UniProt: a worldwide hub of protein knowledge. Nucleic Acids Res. 47, D506–D515.

Van Heuven, S., Pierrot, D., Rae, J. W. B., Lewis, E., and Wallace, D. W. R. (2011). MATLAB program developed for CO_2_ system calculations. ORNL/CDIAC-105b. Carbon Dioxide Information Analysis Center, Oak Ridge National Laboratory, US Department of Energy, Oak Ridge, Tennessee 530.

Venkataraman, Y. R. (2020). Eastern oyster (*Crassostrea virginica*) gonad DNA methylation data and analysis. Available at: https://doi.org/10.6084/m9.figshare.11923479

Venkataraman, Y. R., Spencer, L. H., and Roberts, S. B. (2019). Larval Response to Parental Low pH Exposure in the Pacific Oyster *Crassostrea gigas*. Journal of Shellfish Research 38, 743. doi:10.2983/035.038.0325.

Waldbusser, G. G., Hales, B., Langdon, C. J., Haley, B. A., Schrader, P., Brunner, E. L., et al. (2014). Saturation-state sensitivity of marine bivalve larvae to ocean acidification. Nat. Clim. Chang. 5, 273.

Wang, H.-Q., Tuominen, L. K., and Tsai, C.-J. (2011). SLIM: a sliding linear model for estimating the proportion of true null hypotheses in datasets with dependence structures. Bioinformatics 27, 225–231.

Wright, R. M., Aglyamova, G. V., Meyer, E., and Matz, M. V. (2015). Gene expression associated with white syndromes in a reef building coral, *Acropora hyacinthus*. BMC Genomics 16, 371.

Zhang, X., Li, Q., Kong, L., and Yu, H. (2018). DNA methylation frequency and epigenetic variability of the Pacific oyster *Crassostrea gigas* in relation to the gametogenesis. Fish. Sci. 84, 789–797.

Zhao, L., Liu, L., Liu, B., Liang, J., Lu, Y., and Yang, F. (2019). Antioxidant responses to seawater acidification in an invasive fouling mussel are alleviated by transgenerational acclimation. Aquat. Toxicol. 217, 105331.

Zhao, L., Yang, F., Milano, S., Han, T., Walliser, E. O., and Schöne, B. R. (2018). Transgenerational acclimation to seawater acidification in the Manila clam *Ruditapes philippinarum*: Preferential uptake of metabolic carbon. Sci. Total Environ. 627, 95–103.

